# Biomolecular Condensates Act as Distinct Solvation Environments that Reshape Amino Acid pK_a_ Values

**DOI:** 10.64898/2026.01.11.698869

**Authors:** Shiv Rekhi, Jeetain Mittal

**Affiliations:** Artie McFerrin Department of Chemical Engineering, Texas A&M University, College Station, TX 77843, USA; Department of Chemistry, Texas A&M University, College Station, TX 77843, USA; Interdisciplinary Graduate Program in Genetics and Genomics, Texas A&M University, College Station, TX 77843, USA

## Abstract

Biomolecular condensates create distinct solvation environments in which the ionization equilibria of amino acid side chains may differ markedly from those in bulk aqueous solution. Here, we use all-atom continuous constant pH molecular dynamics simulations to investigate the changes to the pK_a_ values of titratable residues between the coexisting phases of biomolecular condensates. We find that protonated states are favored in the condensate, resulting in the stabilization of charged forms of cationic residues and neutral forms of anionic residues. The effect is consistent across condensates formed by five peptide sequences suggesting that the preference for protonated states is a universal feature of the condensate microenvironment. This highlights that differences in the solvation environments between the coexisting phases play a central role in charge regulation in phase-separating proteins.

---

Biomolecular condensates represent a distinct physicochemical phase of matter in which proteins experience environments that differ fundamentally from bulk aqueous solution^1^. To maintain electroneutrality, charged phase-separating proteins sequester counterions within the dense phase, shaping a unique electrostatic and solvation environment^2-4^. While phase separation is often discussed in terms of concentration-driven organization, the dense phase also imposes altered solvation^5,6^, dielectric response^7,8^, and electrostatic screening^9^ that can directly reshape chemical equilibria. A central but largely unexplored consequence of this altered environment is its impact on charge regulation^10^, which depends sensitively on the solution pH and the pK_a_ values of titratable amino acid side chains. Although recent studies have reported shifts in the pH of the dense phase^11,12^, these changes alone are insufficient to account for the substantial modulation of protein charge expected within condensates^11^, suggesting that the intrinsic pK_a_ values of amino acid side chains may themselves be altered^13,14^ within the condensate microenvironment.

Previous work on charge regulation in disordered proteins^10,15^ and polyelectrolytes^16-19^ has involved the use of implicit solvent or coarse-grained models, which successfully capture pH-dependent effects originating from the electrostatic interactions between monomers. However, solvent reorganization, which requires an explicit representation of solvent molecules, can dominate the charging free energy of a titratable residue^20,21^. Therefore, we use all-atom continuous constant pH molecular dynamics (AA CpHMD) simulations^22^ with explicit solvent^23,24^ to calculate the pK_a_ values of four titratable amino acid side chains (Asp, Glu, His, and Lys), within the dense phase. In AA CpHMD, λ-dynamics^25^ is used to sample protonation states along a pH-dependent free energy surface. The parameters of this surface are calibrated to match the experimental pK_a_ values of the amino acid embedded within a model pentapeptide (ACE-AAXAA-NHE) in solution. By calculating the pK_a_ values of the same pentapeptide placed within a model condensate, we provide an estimate of the pK_a_ shifts due to the condensate microenvironment relative to the dilute (aqueous) phase. The steps to estimate pK_a_ values from AA CpHMD simulations are shown in Figure 1. From our prior work^5^ demonstrating the validity of minimal peptide systems as model condensate systems, we consider the dense phase formed by the minimal Ser-Tyr-Gly-Gln (SYGQ) peptide unit as our model system. The protein and water densities within the condensate were set according to the NMR estimates of the FUS LC condensate^26^.

**Figure 1.**
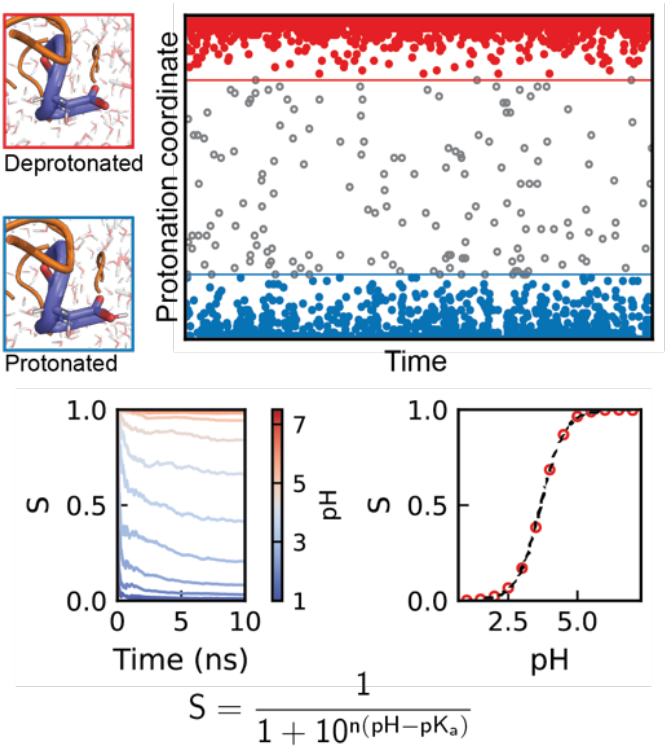
Calculation of pK_a_ values of titratable groups in AA CpHMD simulations. The fraction of deprotonated states (S) is estimated at a user-defined pH condition by tracking the time evolution of the protonation coordinate. Protonated and deprotonated states are assigned based on cutoff values (solid lines) and used to calculate S. Unphysical intermediate points (gray circles) are ignored. Simulations are conducted using a pH-replica exchange approach to calculate S and the resulting S vs. pH curve is fit to the Henderson-Hasselbalch (HH) equation to estimate the pK_a_ of the group along with the Hill coefficient denoted by n.

In the condensate, the titration curves for Asp and Glu are shifted towards more basic conditions, yielding pK_a_ values of 7.36 ± 0.10 and 7.98 ± 0.07 respectively (**Fig. 2, Table 1**). The upshifts of 3.6 units for both Asp and Glu in the condensed phase (**Table 1**) indicate that the protonated neutral form of these amino acids is favored within the dense phase. The difference in the free energy of deprotonation between the phases, given by 2.303*RT* (Δ*pK*_*a*_), shows that deprotonation within the condensed phase is disfavored by 4.98 ± 0.14 kcal/mol and 4.98 ± 0.15 kcal/mol for Asp and Glu respectively.

**Table 1.** pK_a_ values in the dilute and dense phase of the SYGQ condensate and pK_a_ shifts (ΔpK_a_) for Asp, Glu, His, and Lys residues.

| Residue | $pK_a^{dil}$ | $pK_a^{dense}$ | $\Delta pK_a$ |
| --- | --- | --- | --- |
| Asp | $3.73 \pm 0.02$ | $7.36 \pm 0.10$ | $3.63 \pm 0.10$ |
| Glu | $4.35 \pm 0.06$ | $7.98 \pm 0.07$ | $3.63 \pm 0.09$ |
| His | $6.65 \pm 0.10$ | $9.76 \pm 0.03$ | $3.11 \pm 0.10$ |
| Lys | $9.96 \pm 0.06$ | $13.25 \pm 0.06$ | $3.30 \pm 0.09$ |

**Figure 2.**
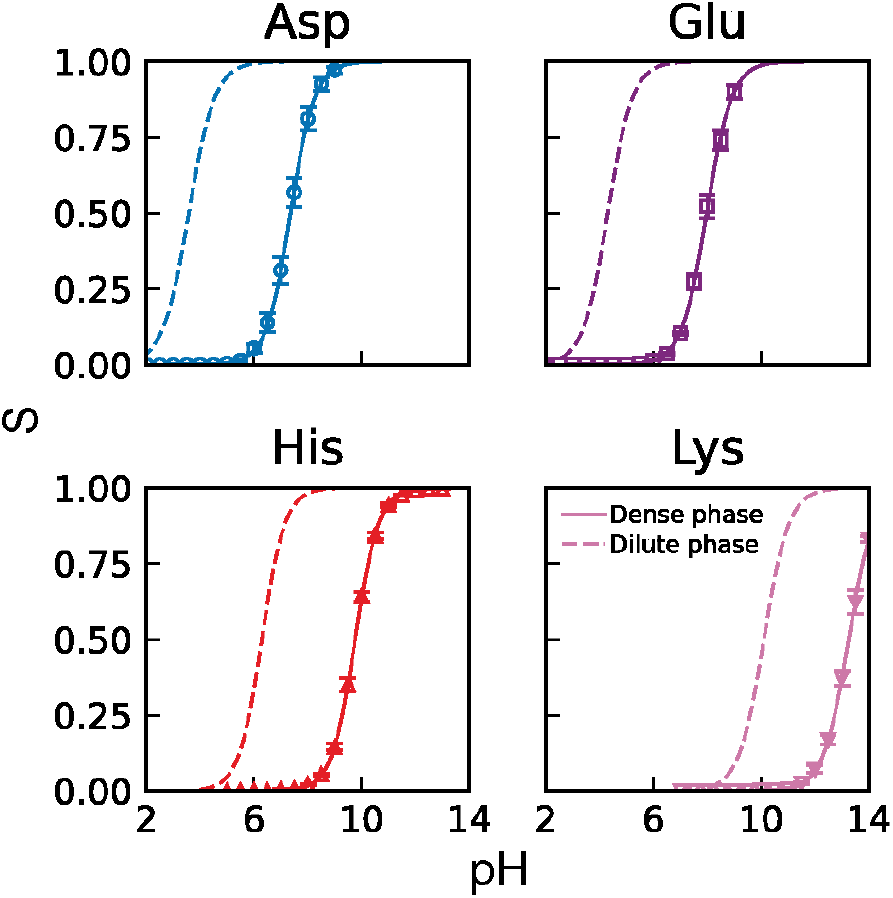
Anionic and cationic residues show elevated pK_a_ values in the dense phase compared to the dilute phase. Titration (S vs. pH) curves for Asp, Glu, His, and Lys residues in the dilute phase (dashed lines) and dense phase (symbols and solid lines). For the dilute phase, only the HH equation fits are shown. For the dense phase, symbols represent the S values estimated from simulation and solid lines show the HH fit to the simulation data. Uncertainties on the simulations are estimated as the SEM over 3 equally spaced 5 ns blocks.

The positive free energy of charging within the condensed phase compared to the dilute phase follows from the expectation of a reduced dielectric constant within the dense phase^7,8^. Similar magnitudes of pK_a_ shifts for Asp have been reported in CpHMD simulations of transmembrane helices^14^. As a baseline estimate of the desolvation penalty originating from different dielectric constants between the phases, we use the Born solvation model^13^. To achieve the charging free energy change observed for Asp, the model requires a dielectric constant of ∼10 in the dense phase, which is significantly lower than the reported values for condensates^7,8^. This indicates that the difference in dielectric constants between phases captures the qualitative preference for protonated states but substantially underestimates the magnitude of the observed pK_a_ shifts, indicating that condensates cannot be treated as simple low-dielectric continua.

Like Asp and Glu, the titration curves for the cationic residues are shifted to higher pH in the condensate and yield pK_a_ values of 9.76 ± 0.03 and 13.25 ± 0.06 for His and Lys (**Fig. 2, Table 1**). The higher pK_a_ values reflect a preference for the positively charged protonated forms within the condensate. Computational^5^ and experimental^27^ estimates of the transfer free energies of charged amino acids and ions^28^ from the dilute to the dense phase report favorable transfer of positively charged species and unfavorable transfer of negatively charged species. The stabilization of protonated states observed here provides mechanistic insight on how charge regulation and transfer free energies jointly dictate the solvation of titratable molecules in condensates. Shifting the protonation equilibria of anions to favor neutral forms allows for their incorporation within the condensate microenvironment despite the unfavorable transfer of their charged forms. Conversely, charged forms of cations are stabilized, reflecting their thermodynamically favored incorporation into the dense phase.

The difference in the free energy of charging between phases for cationic amino acids is favorable within the condensed phase with Lys (-4.52 ± 0.14 kcal/mol) being marginally more negative than His (-4.26 ± 0.12 kcal/mol). This contrasts with the expectation from the Born model which predicts unfavorable charging free energy in a medium of lower dielectric than water. This implies that the dielectric-dependent desolvation term must be compensated for by favorable interactions with the environment^20^. Here, these compensating “background” interactions arise from surrounding permanent dipoles and charged molecules. However, isolating their individual contributions remains challenging. We therefore exploit sequence-dependent variations in condensate composition to probe the balance between desolvation and background interactions by comparing pK_a_ shifts of titratable residues in condensates formed by different peptide sequences.

The additional sequences investigated were: (i) a variant of the elastin-like polypeptide (ELP, APGVG), (ii) the resilin-like polypeptide (RLP, GRGDSPYS), (iii) a positively charged RLP variant (RLP^+^, GRGNSPYS), and (iv) a negatively charged RLP variant (RLP^-^, GQGDSPYS). We find that between all systems the qualitative trend of favorable protonation within the condensed phase is preserved (**Fig. 3, Table 2, Fig. S1**). Moreover, the magnitude of pK_a_ shifts for all residues are relatively consistent across all systems investigated in this work, indicating that the observed pK_a_ shifts are only weakly sequence-dependent and instead primarily governed by the shared solvation characteristics of the condensed phase (**Fig. 3, Table 2**).

**Table 2.** pK_a_ values and pK_a_ shifts with respect to the dilute phase (ΔpK_a_) for Asp, Glu, His, and Lys residues in the APGVG, RLP, RLP^+^, and RLP^-^ condensates.

| Residue | $pK_a^{APGVG}$ | $\Delta pK_a^{APGVG}$ | $pK_a^{RLP}$ | $\Delta pK_a^{RLP}$ | $pK_a^{RLP^+}$ | $\Delta pK_a^{RLP^+}$ | $pK_a^{RLP^-}$ | $\Delta pK_a^{RLP^-}$ |
| --- | --- | --- | --- | --- | --- | --- | --- | --- |
| Asp | 8.07±0.16 | 4.33±0.16 | 7.44±0.08 | 3.71±0.09 | 6.87±0.09 | 3.13±0.10 | 7.28±0.16 | 3.54±0.16 |
| Glu | 8.80±0.06 | 4.45±0.08 | 7.93±0.04 | 3.57±0.07 | 7.68±0.10 | 3.33±0.12 | 7.32±0.02 | 2.97±0.06 |
| His | 10.60±0.04 | 3.95±0.10 | 10.38±0.08 | 3.73±0.13 | 9.89±0.01 | 3.24±0.10 | 9.76±0.02 | 3.11±0.10 |
| Lys | 13.99±0.08 | 4.03±0.10 | 13.54±0.05 | 3.58±0.08 | 13.11±0.11 | 3.16±0.12 | 13.27±0.08 | 3.31±0.10 |

**Figure 3.**
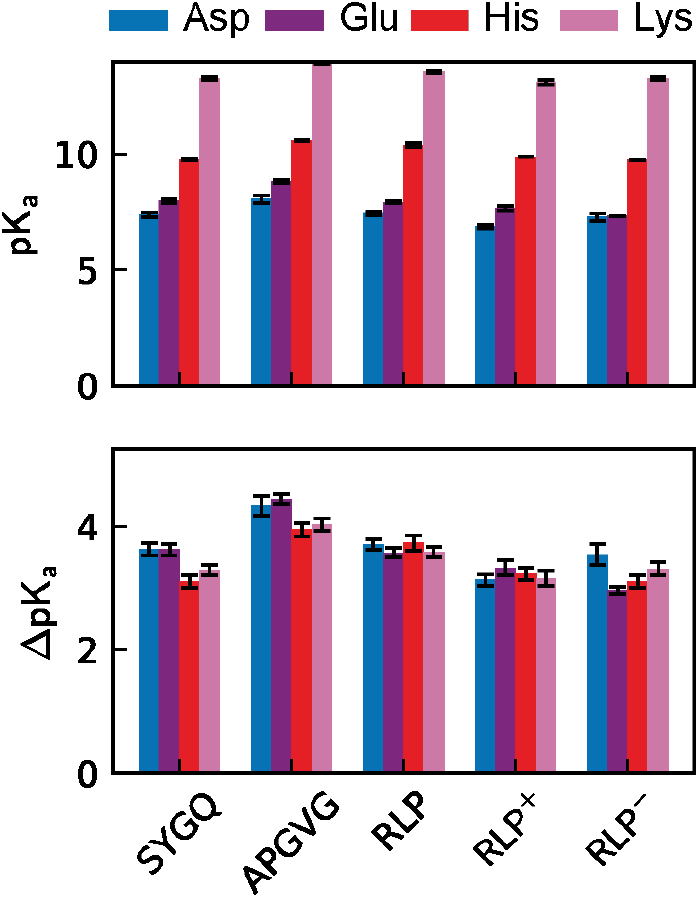
Significant stabilization of protonated states is observed across 5 condensate systems. pK_a_ values (upper panel) and pK_a_ shifts with respect to the dilute phase (lower panel) for Asp, Glu, His, and Lys in condensates formed by the SYGQ, APGVG, RLP, RLP^+^, and RLP^-^ peptide sequences. pK_a_ and ΔpK_a_ values for the SYGQ condensate are identical to those in Table 1. Uncertainties for all residues and condensate systems are estimated as the SEM over 3 equally spaced 5 ns blocks. The consistency of the pK_a_ shifts across chemically distinct condensates highlights a general solvation-driven effect of the dense phase rather than sequence-specific interactions.

Compared to SYGQ, in APGVG we see higher upshifts for both the anionic and cationic amino acids (**Fig. 3, Table 2**). This suggests that the absence of polar side chains in the peptide sequence leads to further destabilization of the ionized negative form for anions and stabilization of the positive form for cations. Between SYGQ and RLP, we observe smaller differences in the pK_a_ shifts (**Fig. 3, Table 2**), except for His where in RLP, the charged form is marginally stabilized, implying that the addition of charged residues has a minimal effect. Perhaps most surprisingly, the preference for protonated states within the condensate compared to the dilute phase persists even in RLP^+^ and RLP^-^ (**Fig. 3, Table 2**). Relative to SYGQ, in RLP^+^ we see that the associative charging behavior seen in complexation of polyelectrolytes^17-19^, where pK_a_ values shift to minimize charge repulsion and maximize charge interactions, is qualitatively reflected in the mean values of all amino acids. However, the effect is less pronounced than expected (**Fig. 3, Table 2**). In the case of RLP^-^, we see that the expectation of upshifted pK_a_ values with respect to SYGQ is not reflected even in the mean values (**Fig. 3, Table 2**).

The range of pK_a_ values observed in the different systems for Asp (6.86–8.07), Glu (7.32– 8.80) and His (9.75–10.60) indicates that small pH shifts in the dense phase can significantly alter the net charge on proteins. Therefore, we compute the net-charge-per-residue (NCPR) profiles of four charge-rich IDRs (hnRNPA1-LCD, DDX4-NTD, LAF1-RGG, RLP-R-to-K) as a function of pH (**Fig. 4**). Within biologically relevant range of pH values of 4.5–8, we see that all the proteins have a positive NCPR (**Fig. 4**). Accounting for the pK_a_ shifts in the condensates also leads to a higher magnitude of NCPR across the pH range than expected from the model pK_a_ values (**Fig. 4**). From this, we anticipate that IDRs with high content of cationic or anionic residues will have more basic microenvironments than predicted from their isoelectric point (pI). Estimates of the dense phase pH of condensates formed by some sequences in the literature lend some support to this hypothesis. For instance, charge-rich, but net-neutral RLP condensates^3^ have a pH of ∼8 (pI = 6.43) and net negatively charged PGL3 condensates^12^ have a pH of ∼6 (pI = 5.1). But for FUS^12^, the experimentally measured pH of ∼8.5, is more acidic than expected from its pI (9.4). To resolve this, further investigation is required to understand the conformational dependence of the pK_a_ values of the titratable residues and the interplay between ion partitioning, dense phase pH, and protein net charge in determining the internal electrostatic environment of condensates^9^.

**Figure 4.**
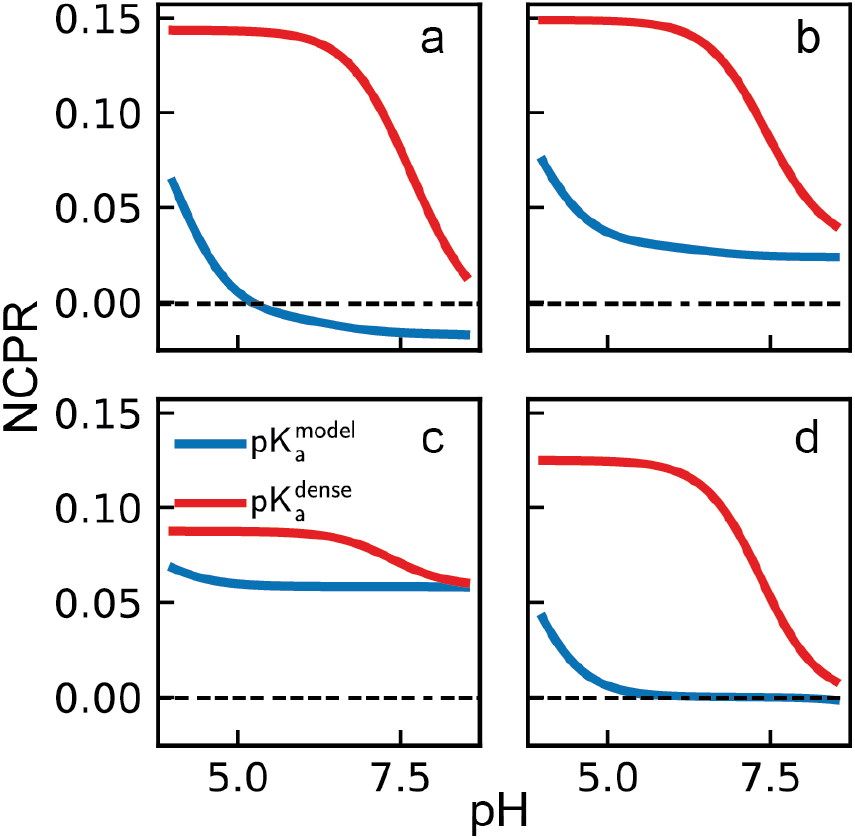
pK_a_ shifts in the dense phase lead to positively charged proteins across a wider range of pH values. Net charge per residue (NCPR) as a function of solution pH for (a) DDX4-NTD, (b) LAF1-RGG, (c) A1-LCD, and (d) an RLP variant with all R mutated to K. Blue lines show profiles when using model compound (aqueous) pK_a_ values show in Table S1. Red lines show profiles calculated using the dense phase pK_a_ values shown in Table 1. The charge on Arg is always taken as +1 within the range of pH values considered. Cys, Tyr, and the backbone titratable groups are all considered neutral.

In summary, our results demonstrate that biomolecular condensates act as chemically distinct solvation environments that strongly stabilize protonated states of cationic residues and neutral states of anionic residues, leading to large and systematic pK_a_ shifts. The consistency of this effect across multiple condensate-forming sequences suggests that charge asymmetry^5,27,29^ is a general feature of condensate solvation rather than a sequence-specific anomaly. By directly quantifying how the condensate microenvironment reshapes ionization equilibria, this work reveals an unrecognized mechanism of charge regulation in phase-separated systems. These findings have broad implications for understanding how sequence composition, electrostatics, and environmental coupling jointly determine the emergent chemical properties of biomolecular condensates^9,30^.

## Acknowledgements

This work was supported by the National Institute of General Medical Sciences of the National Institute of Health (R35GM153388). We thank Dr. Busra Ozguney (Texas A&M University) for running tests of CpHMD simulations within the condensate and aiding in the preparation of some of the systems used in this work. We gratefully acknowledge the Texas A&M High Performance Research Computing for providing the computational resources for this work.

