## Supplementary Information for "Biomolecular Condensates Act as Distinct Solvation Environments that Reshape Amino Acid pK_a_ Values"

### Supporting Text

### Methods

*Preparation and equilibration of condensate systems*

An initial pdb file for a single chain of the capped peptide sequence (ACE-SYGQ-NHE) was prepared using the LEAP module in AmberTools[1]. Following this, 80 chains were inserted in a 5 nm side cubic simulation box using PACKMOL[2], corresponding to a peptide concentration of ~500 mg/ml. The system was then solvated and 11 $Na^+$ and $Cl^-$ ions were added to achieve a salt concentration of 150 mM. The system was modeled using the AMBERFF14SB[3] protein forcefield in combination with the TIP3P[4] water model. Cufix corrections were applied to improve the electrostatic interactions in the forcefield[5,6]. Hydrogen masses were repartitioned by a factor of 1.5 to allow simulations with a time step of 4 fs[7].

The system was then minimized for 3000 steps (1000 steps steepest descent, followed by conjugate gradient) with the proteins harmonically restrained with a force constant of 50 kcal/mol/Å$^2$. Following this, a series of equilibration steps were carried out to gradually release the restraints on the protein, increase the temperature to the target value of 300K, and increase the time step to 4fs. First, we simulated the system for 5000 steps with a time step of 0.5 fs at 100K and 1 bar using the Langevin thermostat and Berendsen barostat[8]. Restraints were maintained on the protein with the same force constant of 50 kcal/mol/Å$^2$. In the second step, the time step was increased to 1 fs with the temperature maintained at 100K, pressure at 1 bar, and protein restraints maintained at 50 kcal/mol/Å$^2$. The third step involved heating the system from 100K to 300K, increasing the time step to 2 fs and reducing the force constant on the restraints to 25 kcal/mol/Å$^2$. In the fourth step, the system was simulated for 10000 steps at a temperature of 300K with a time step of 4 fs. The pressure coupling algorithm was changed to the Monte Carlo barostat in this step. Following the four equilibration steps, the system was run on the GPU using the pmemd.cuda executable in AMBER24[9] for 20000 steps in the NPT ensemble with temperature 300K, pressure 1 bar, and a time step of 4 fs using the Langevin thermostat and Monte Carlo barostat. This simulation was then continued for 250000000 steps for a total run time of 1 μs. Coordinates were written to the trajectory file every 100 ps leading to 10000 frames from which densities were calculated using modules in MDAnalysis. The first 200 ns were discarded from the analysis as equilibration.

The density profiles from the above simulation with the AMBER14SB+TIP3P forcefields showed that proteins clustered within the simulation box (**Fig. S2**). To prevent the collapse of the protein chains into a phase of higher concentration within the simulation box, we scaled the protein-water Lennard-Jones (LJ) interactions by a factor of 1.1[10]. We used the same initial solvated configuration of peptide chains and used the Parmed module in AmberTools[1] to apply the scaling of LJ interactions. The system was equilibrated following the same steps as detailed above and the density of protein within the simulation box was calculated. The density profile showed an improvement compared to the unscaled AMBER14SB+TIP3P water model. Therefore, the last frame from this simulation was used as the structure for inserting the model pentapeptides used to estimate the $pK_a$ values in the condensed phase.

The additional condensate systems (APGVG, RLP, $RLP^+$, and $RLP^-$) were prepared following the same steps. 80 chains of APGVG and 40 chains of RLP, $RLP^+$, and $RLP^-$ were used to achieve the desired peptide concentrations in the simulation box. For the $RLP^+$ and $RLP^-$ systems, counterions were added to ensure neutrality within the simulation box. All systems were modeled using the scaled AMBER14SB protein forcefield in combination with the TIP3P water model.

*The all-atom CpHMD simulation method*

All-atom continuous constant pH molecular dynamics (AA CpHMD) simulations sample the time evolution of protonation states of a titratable residue using $\lambda$-dynamics[11,12]. In AA CpHMD the titration coordinate ($\lambda$ for single site titration and $\lambda$ and x in double site titration), is propagated using an extended Hamiltonian. We point readers to recent reviews and research articles for comprehensive explanations of the method[13,14].

The pH-dependence of the titration coordinate is included through biasing potentials that ensure correct pH-dependent sampling. The parameters for these biasing potentials are calibrated such that the $pK_a$ of the titratable residue embedded within a model compound in solution matches the experimentally determined $pK_a$. We used the parameter files for the AMBER14SB forcefield and TIP3P water model combination deposited on the JanaShenLab GitLab page in our simulations[11,15]. As the Lys parameters were not available for AA CpHMD simulations with the AMBER14SB forcefield, we generated them following the instructions provided in a recent review[16].

The calculation of $pK_a$ values involves running the system at multiple solution pH conditions and calculating the fraction of deprotonated states denoted by S. These values are then fitted to the Henderson-Hasselbalch equation shown below to calculate the $pK_a$ values and the Hill coefficient (n) of the titratable group.

$$S = \frac{1}{1 + 10^{n(pK_a - pH)}}.$$

Values of n<1 and n>1 indicate anti-cooperativity and cooperativity respectively[12]. S values are calculated considering λ<0.2 and λ>0.8 as protonated and deprotonated respectively. To improve the sampling of the titration coordinates, we used the asynchronous pH-replica exchange sampling scheme[17]. Details of the pH ranges and runtimes are provided below for the dilute phase and dense phase simulations.

*CpHMD simulations of model pentapeptides in the dilute phase: Effect of scaled interactions*

Pdb files of the blocked model pentapeptides containing the titratable amino acids were generated using the LEAP module in AmberTools. Modifications were made to the residue names for the titratable groups in the pdb file to make them compatible with the CpHMD input parameter files (ASP to AS2, GLU to GL2, HIS to HID). The forcefield modification (frcmod) files required to define the titratable residue parameters files for the CpHMD simulations were downloaded from the JanaShenLab GitLab page[15]. Using the LEAP module, the pentapeptide pdb file was placed in a cubic water box (TIP3P) of side 5 nm, counterions (calculated based on charge state of residue at neutral pH) and ions to maintain a concentration of 150mM were added, and the AMBER parameter and coordinate files were generated. Hydrogen masses were repartitioned (1.5 Da) to enable simulations with a time step of 4 fs. The systems were then minimized and equilibrated following the same procedure as for the preparation of the condensed phases detailed above. Following the equilibration steps, the system was run for 2.5 ns with pH-dependent sampling applied at neutral pH to allow the titration coordinates to equilibrate. The last frame from this simulation was extracted and used as the initial structure for the replica exchange CpHMD production simulations.

We ran asynchronous pH-replica exchange simulations using the codes provided by the Shen group[15]. The range of pH values for the different amino acids were, 1-7.5 in steps of 0.5 for Asp and Glu, 3-9 in steps of 0.5 for His, and 8-13 in steps of 0.5 for Lys. The total runtime for each window was 12.5 ns, with swaps attempted every 2.5 ps. The values of the titration coordinates were stored every 1.25 ps for analysis.

We ignored the first 3.12 ns of sampling as equilibration and block averaged the fraction of deprotonated states (S) over 3 equally spaced blocks of the remainder of the trajectory. $pK_a$ values were estimated for each of the blocks by fitting the HH equation. Final values are reported as the mean of the $pK_a$ values of the 3 blocks and uncertainties are reported as the SEM over three 3 blocks.

The $pK_a$ values in AA CpHMD are dependent on the parameters of the biasing potentials applied in the extended Hamiltonian. These parameters differ between forcefields. To verify whether the scaling of protein-water interactions requires a reparameterization of the biasing potentials, we compared the $pK_a$ values of the four titratable residues calculated using the scaled and unscaled AMBER14SB forcefields. The $pK_a$ values between the two forcefields were within statistical uncertainty of each other for all the residues (**Fig. S3, Table S1**). Thus, we did not make any changes to the pH-dependent sampling parameter file.

*CpHMD simulations of the titratable residues within the condensed phases*

We extracted the last frame of the condensate from the 1 μs simulations and used the GROMACS[18] command *gmx-insert* to place a copy of the blocked residue within the simulation box. We took the resulting file in pdb format and used LEAP with the CpHMD frcmod files to prepare the parameter files for CpHMD simulations. The systems were equilibrated and minimized following the same steps as the dilute phase calculations.

Within the condensates, to account for the $pK_a$ shifts, we used a spacing of 0.5 units and pH ranges of pH-replica exchange simulations. We ran the simulations for a total runtime of 20 ns per window, attempting swaps every 4 ps and the values of the titration coordinates were stored every 2 ps. We saw that for all the amino acids in the SYGQ condensate, the replicas exchanged freely (**Fig. S4**) suggesting adequate sampling with the chosen swap frequencies. Within the SYGQ condensate, it took ~3-5 ns for the running average of the fraction of deprotonated residues for all residues to flatten (**Fig. S5**). A similar timescale for the plateauing of the running average of S was observed in other systems as well (**Fig. S6**). Therefore, we discarded the first 5 ns of sampling and performed our analysis on the remainder of the trajectory for all amino acids in all condensate systems. $pK_a$ values and uncertainties are reported using the same protocol as detailed for the dilute phase with the only change being the use of 3 blocks of 5 ns each.

In AA CpHMD simulations carried out in AMBER without titratable water, there is a net charge within the simulation box due to the insertion of a counterion in the case of Asp, Glu, and His. This is due to the convention of adding counterions in the initial structure assuming charge states at neutral pH, though the simulations are always started from the protonated states. This leads to the application of a neutralizing background charge when using PME electrostatics which can artificially stabilize one charge form over another[19]. This effect of the neutralizing charge is expected to reduce with increasing box size. To test the role of this artefact in sampling, we calculated the $pK_a$ value of Asp within an SYGQ condensate prepared in a larger simulation box of side 7.5 nm. The system was prepared and equilibrated following the same steps as for the other systems. The final frame from this simulation was extracted, the model pentapeptide was inserted, and the simulations to calculate the $pK_a$ were conducted following the steps and run parameters detailed above. Our test revealed that the mean $pK_a$ value of Asp reduced by ~0.20 within the larger box compared to our 5 nm side system (**Fig. S7**). This suggests that while there is an effect of the size of simulation cell, the error introduced by the finite-size of the box is much smaller than the reported $pK_a$ shifts.

*Computation of net-charge-per-residue profiles*

We computed the net-charge-per-residue for the four IDP sequences using the $pK_a$ values in the dense phase and the model pentapeptide in solution. The experimentally determined $pK_a$ values for Asp, Glu, His, and Lys, shown in Table S1, were used as the model $pK_a$ values, while the $pK_a$ values of the residues in the SYGQ condensate shown in Table 1 of the main text were used as the dense phase $pK_a$ values. We considered Arg to always be charged within the pH range investigated. We did not consider the possible changes in charge state arising from other titratable groups such as the backbone, tyrosine, and cysteine in our calculations. The range of pH values designated as biologically relevant spans from 4.5 to 8 representative of the pH of the lysozyme (4.5-5.0) and the mitochondrial matrix and some membraneless compartments (~8).

**Supporting Figures**

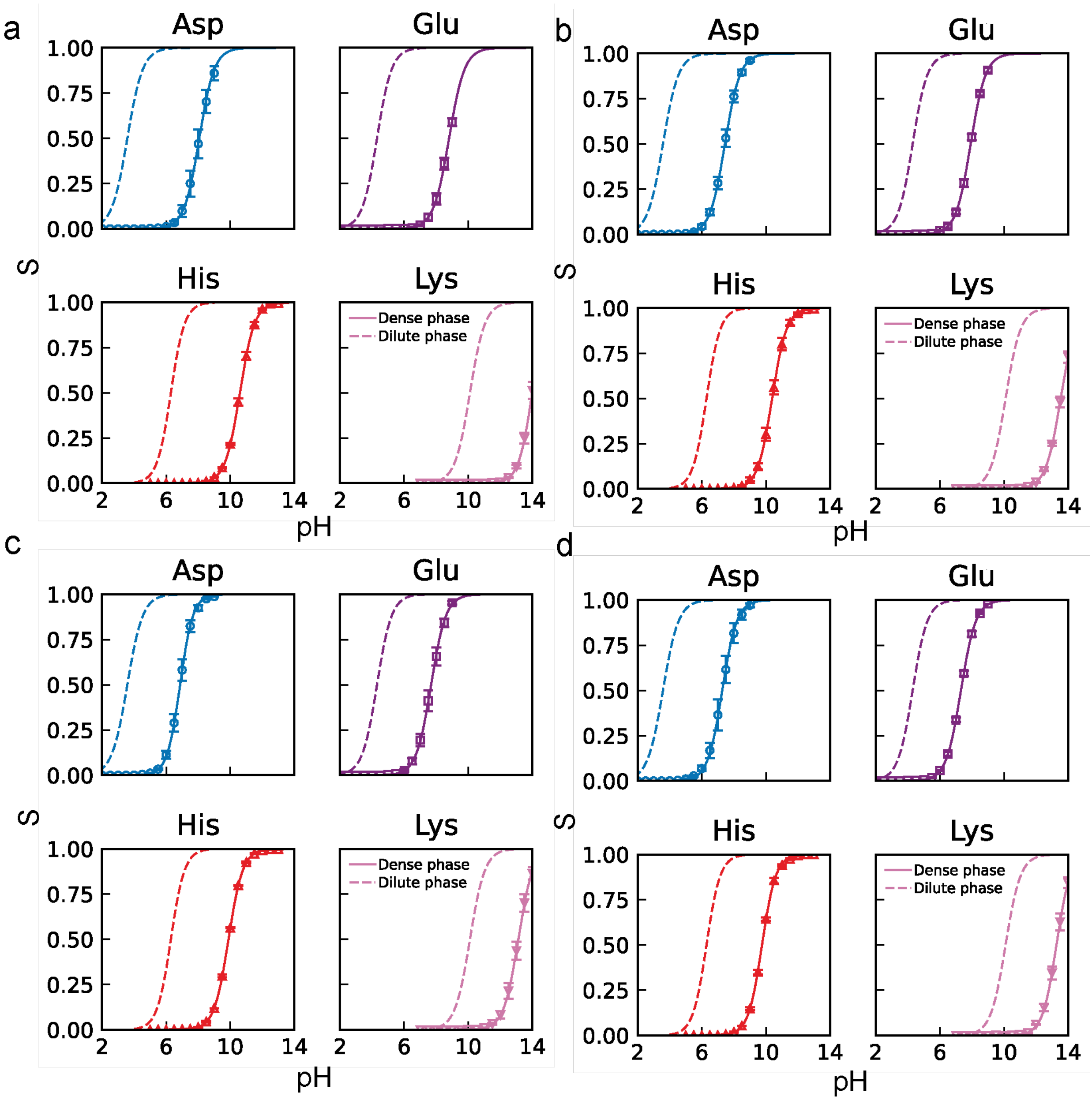


***Figure S1.*** *Titration curves for Asp, Glu, His, and Lys in dilute phase (dashed lines) and dense phase (symbols and solid lines) of the (a) APGVG, (b) RLP, (c) RLP$^{+}$, and (d) RLP$^{-}$ condensates. For the dilute phase, only the HH equation fits are shown. For the dense phase, symbols represent the S values estimated from simulation and solid lines show the HH fit to the simulation data. Uncertainties on the S values from simulation are estimated as the SEM from 3 equally spaced 5ns blocks.*

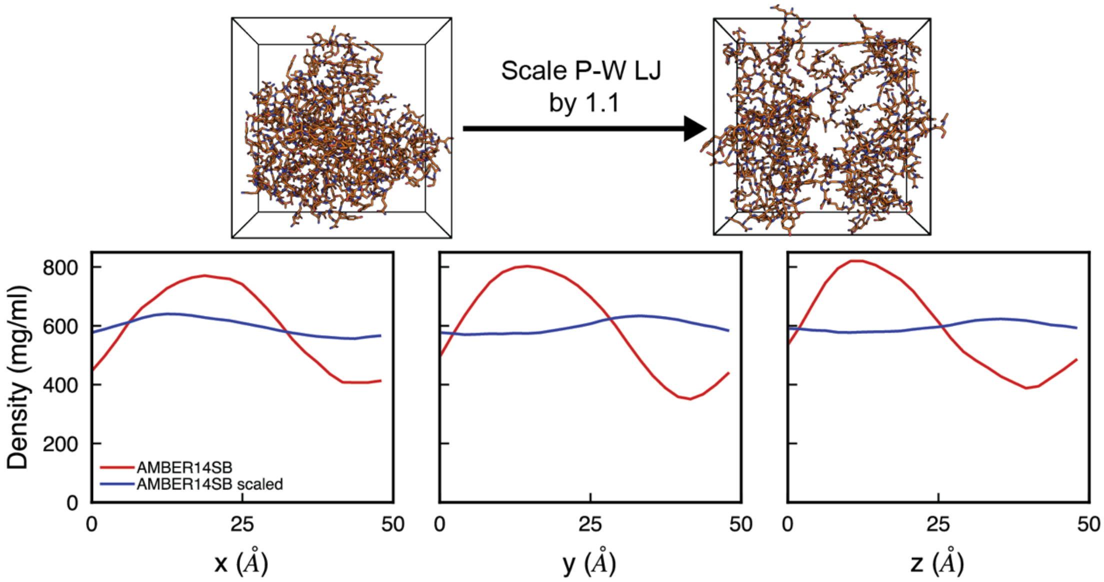


***Figure S2.*** *Comparison of the density of protein in the x, y, and z dimensions of the simulation box for the AMBER14SB forcefield (red) and a variant of the forcefield AMBER14SB scaled (blue), where Lennard-Jones interactions between protein atoms and water oxygens are scaled by 1.1 times. Simulation snapshots above show the protein configurations at the end of a 1μs long NPT simulation. Water molecules and salt ions are hidden for clarity.*

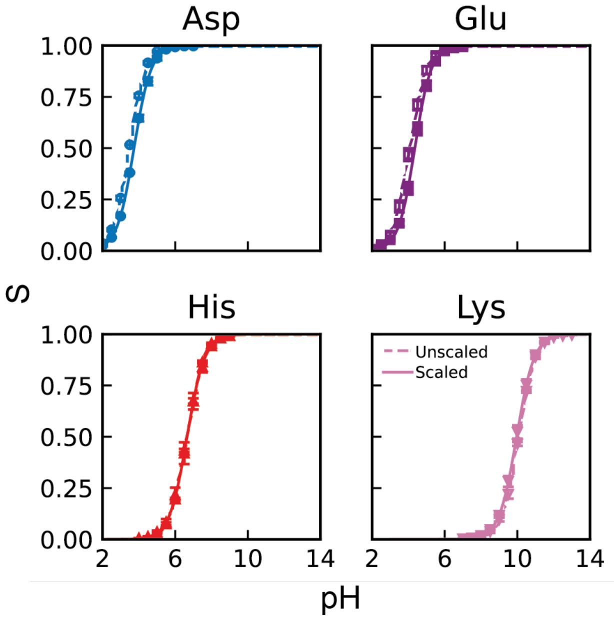


***Figure S3.*** *Comparison of titration curves for Asp, Glu, His, and Lys in the dilute phase using the AMBER14SB protein forcefield (unscaled) and a variant of AMBER14SB where protein–water oxygen Lennard-Jones interactions are scaled by a factor of 1.1 (scaled). p$K_a$ values calculated from the titration curves are tabulated in Table S1.*

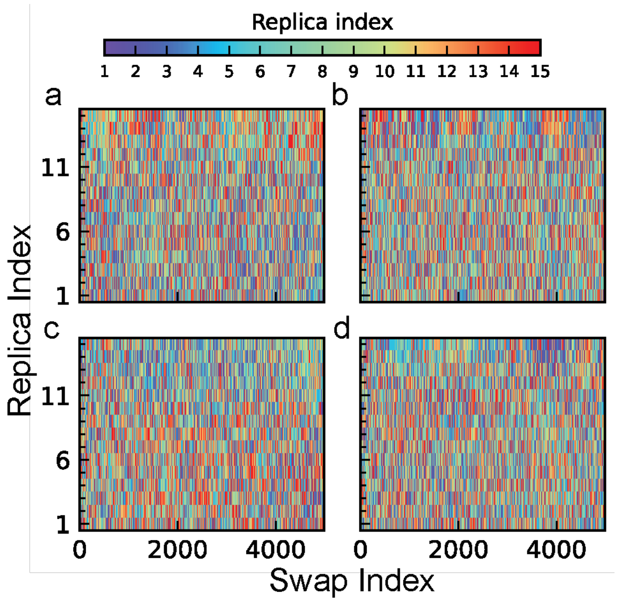


***Figure S4.*** *Replica walks for (a) Asp, (b) Glu, (c) His, and (d) Lys in the SYGQ condensate. 500 swaps are shown, and swaps are attempted every 0.4ns.*

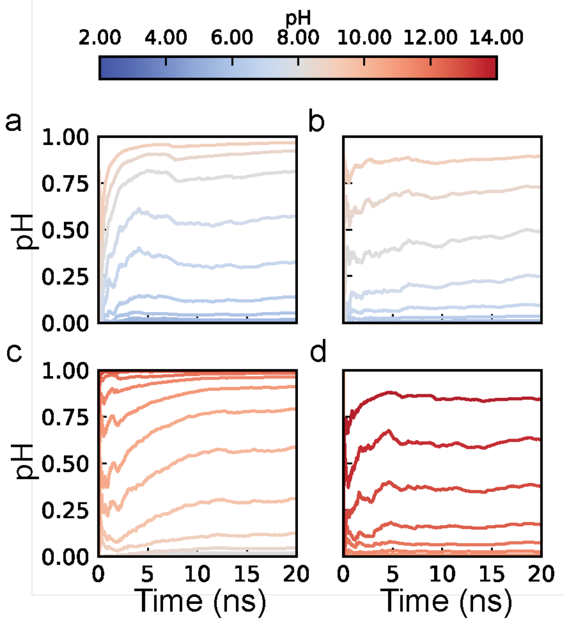


***Figure S5.*** *Running average of deprotonated fraction (S) with time at different solution pH values for the (a) Asp, (b) Glu, (c) His, and (d) Lys systems within the SYGQ condensate.*

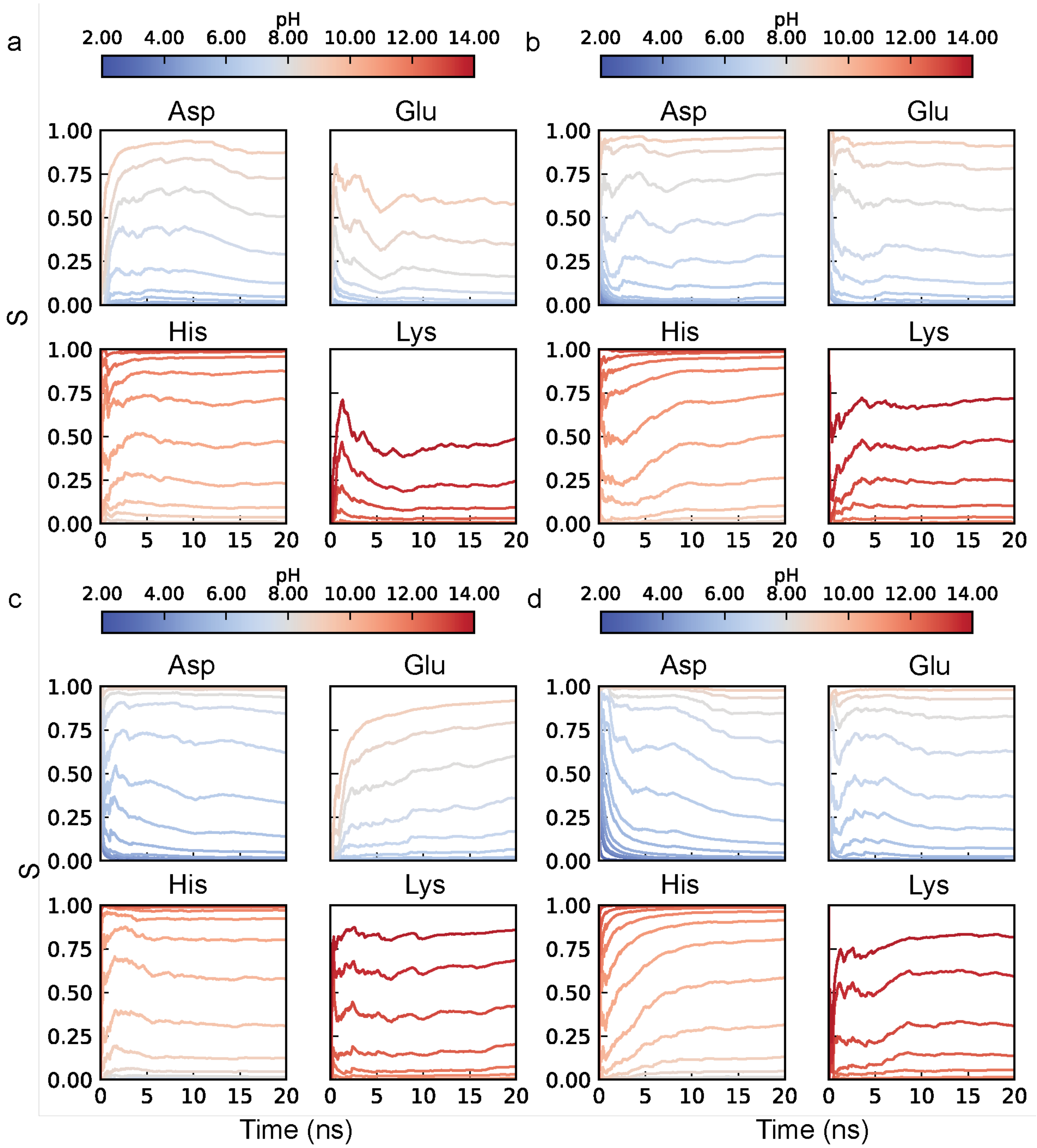


***Figure S6.*** *Running average of deprotonated fraction (S) with time at different solution pH values for Asp, Glu, His, and Lys in the (a) APGVG, (b) RLP, (c) RLP⁺, and (d) RLP⁻ condensates.*

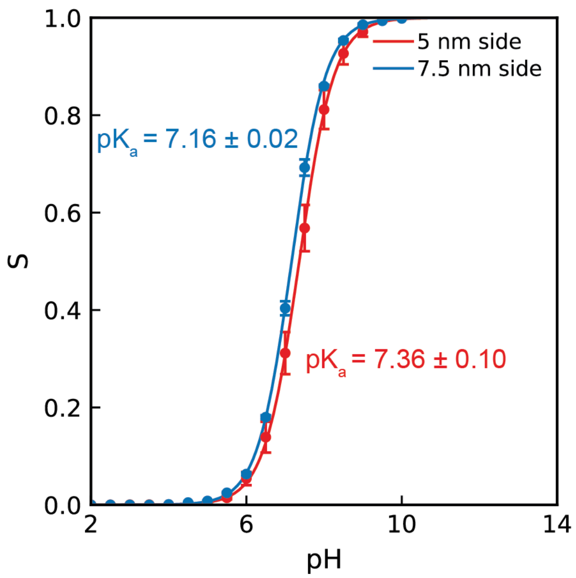


***Figure S7.*** *Comparison of p$K_a$ values of Asp estimated within the SYGQ condensate prepared in 5 nm side cubic box and a 7.5 nm cubic side box.*

### Supporting Tables

| Residue | p$K_a$ (Experiments) | p$K_a$ (AMBER14SB) | p$K_a$ (AMBER14SB scaled) |
|---|---|---|---|
| Asp | 3.70 | 3.48±0.01 | 3.73±0.02 |
| Glu | 4.20 | 4.08±0.06 | 4.35±0.06 |
| His | 6.50 | 6.64±0.01 | 6.65±0.10 |
| Lys | 10.40 | 10.06±0.01 | 9.96±0.06 |

***Table S1.*** *Comparison of p$K_a$ values of Asp, Glu, His, and Lys embedded in the model pentapeptide (ACE-AAXAA-NHE) using the scaled and unscaled variants of the AMBER14SB forcefield. Experimental values are provided as a reference.*
